# Mitochondrial genome recovery by ATFS-1 is essential for development following starvation

**DOI:** 10.1101/2022.05.19.492689

**Authors:** Nandhitha Uma Naresh, Tomer Shpilka, Qiyuan Yang, Yunguang Du, Cole M. Haynes

## Abstract

Nutrient availability regulates the *C. elegans* life cycle as well as mitochondrial physiology. Food deprivation significantly reduces mitochondrial genome (mtDNA) number and leads to aging-related phenotypes. Here, we demonstrate that the bZIP protein ATFS-1, a mediator of the mitochondrial unfolded protein response (UPR^mt^), is required to promote growth and establish a functional germline following prolonged starvation. Surprisingly, we found that the recovery of mtDNA copy number and development following starvation required mitochondrial-localized ATFS-1 but not its nuclear transcription activity. Lastly, we found that the insulin-like receptor DAF-2, functions upstream of ATFS-1 to modulate mtDNA content. We demonstrate that reducing DAF-2 activity represses ATFS-1 nuclear function while causing an increase in mtDNA content partly mediated by mitochondrial-localized ATFS-1. Combined, our data indicate the importance of the UPR^mt^ in recovering mitochondrial mass and suggests that *atfs-1*-dependent mtDNA replication precedes mitochondrial network expansion following starvation.

## INTRODUCTION

The mitochondrial unfolded protein response (UPR^mt^) is a signaling pathway mediated by the bZip protein ATFS-1 that promotes mitochondrial network expansion by regulating transcription of mitochondrial biogenesis genes (Shpilka et al., 2021). The UPR^mt^ transcriptional response has been best studied in the context of mitochondrial perturbations that slow worm development such as mutations in genes encoding OXPHOS proteins, mtDNA heteroplasmy or pathogen infection (Deng et al., 2019; Durieux et al., 2011; Gitschlag et al., 2016; Lin et al., 2016; Moullan et al., 2015; Yoneda et al., 2004). During mitochondrial stress or dysfunction, a fraction of ATFS-1 accumulates in the nucleus via its nuclear localization sequence (NLS) where it activates a transcription program to recover mitochondrial function (Nargund et al., 2015). ATFS-1 also accumulates in dysfunctional mitochondria via its mitochondrial targeting sequence (MTS), where it binds mtDNA and promotes replication (Nargund et al., 2015; Yang et al., 2022). The regulation of both the nuclear transcriptional response as well as mtDNA replication by ATFS-1 during mitochondrial stress is consistent with the UPR^mt^ promoting mitochondrial function by coordinating mitochondria-to-nuclear communication (Figure 1A). However, little is known regarding the role of ATFS-1 in coordinating such a response when mitochondrial dysfunction is not induced by exogenous stressors or deleterious mutations.

**Figure 1.**
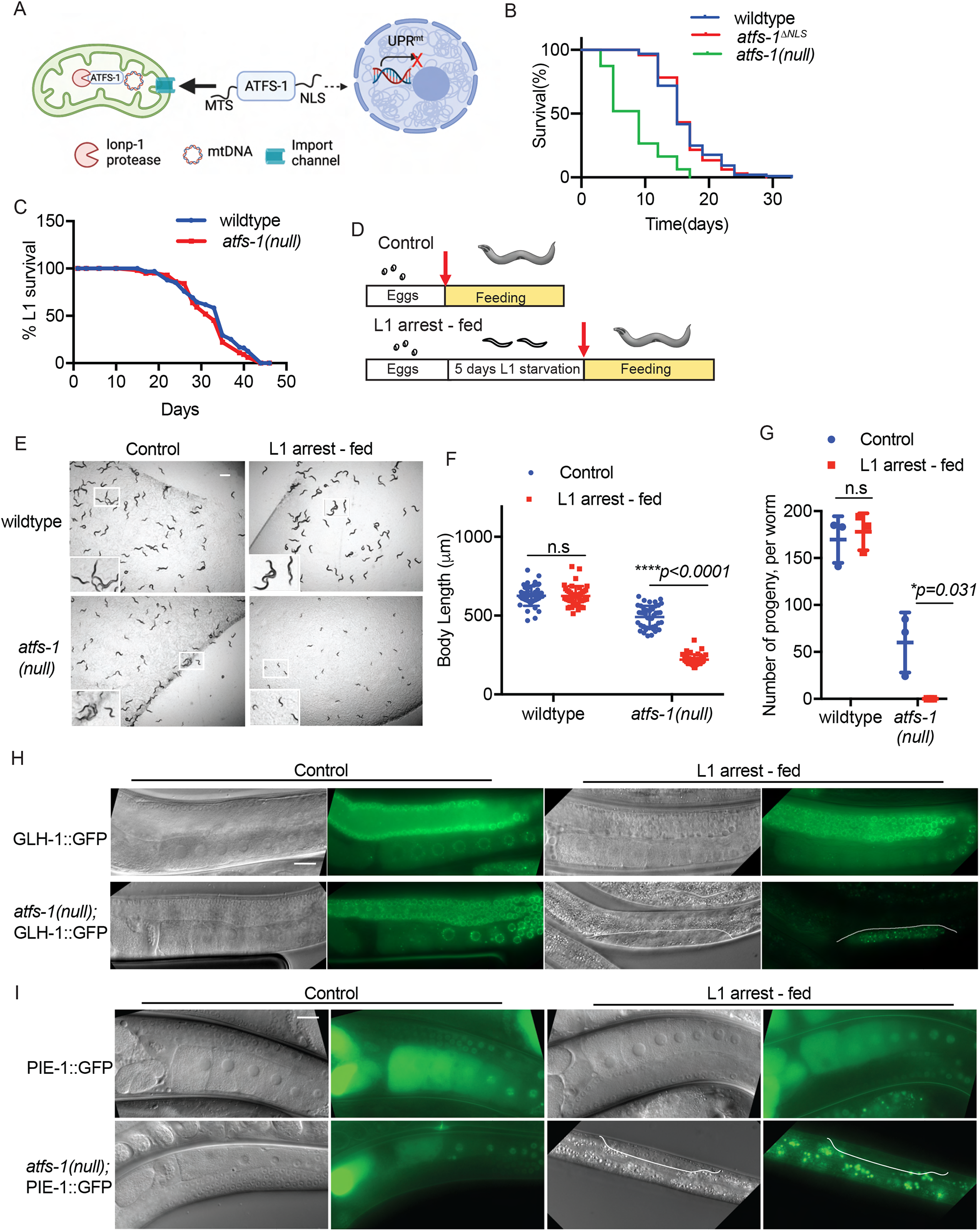
ATFS-1 is required for development following prolonged L1 arrest. A. Schematic illustrating ATFS-1 mediated UPR^mt^ signaling. B. Lifespan analysis comparing wildtype, *atfs-1(null)* and *atfs-1*Δ^*NLS*^ worms (Statistical analyses from three independent repeats are reported in Table S1). C. Lifespan analysis comparing wildtype and *atfs-1(null)* worms during L1 arrest (Statistical analyses from three independent repeats are reported in Table S1). D. Schematic depicting experimental protocol comparing worms that were directly hatched in the presence of bacteria (control) and worms that were maintained as arrested L1s for 5 days and then transferred to plates seeded with bacteria (L1 arrest-fed). E. Images of wildtype and *atfs-1(null)* worms in control and L1 arrest-fed conditions obtained when control worms reached the L4 stage (Scale bar, 0.5 mm). F. Quantification of body lengths comparing wildtype and *atfs-1(null)* worms in control and L1 arrest-fed conditions (*n = 3* ± SD, *\*\*\*\*p<0*.*0001*, Student’ s t-test). G. Brood size quantification of wildtype and *atfs-1(null)* worms in control and L1 arrest-fed conditions (*n = 3* ± SD, *\*p = 0*.*031*, Student’ s t-test). H. Images of wildtype and *atfs-1(null)* worms expressing GLH-1::GFP under control or L1 arrest-fed conditions obtained on day 1 of adulthood (Scale bar, 0.02 mm). I. Images of wildtype and *atfs-1(null)* worms expressing PIE-1::GFP under control or L1 arrest-fed conditions obtained on day 1 of adulthood (Scale bar, 0.02 mm).

Diverse species, including vertebrates, adapt cellular responses to survive adverse environmental conditions including prolonged starvation and extreme seasonal changes such as drought (Hand et al., 2016; Hu and Brunet, 2018; Hu et al., 2020; Van Gilst, 2020). A relatively common adaptation is when the animals enter a diapause state upon encountering severe stress where growth is suspended but basal metabolic activities are maintained. The nematode *C. elegans* can undergo developmental arrest during starvation and resume development once food is encountered (Baugh, 2013; Carranza-Garcia and Navarro, 2020). When worms hatch in the absence of food, they remain arrested at the first larval stage, which is known as L1 diapause. Developmental arrest is accompanied by an increase in stress resistance and alterations in metabolic signaling pathways including insulin signaling, TGF-β signaling, AMP-activated protein kinase, and fatty acid metabolism (Baugh and Sternberg, 2006; Demoinet et al., 2017; Kang et al., 2007; Kaplan et al., 2015; Lee and Ashrafi, 2008). To survive during L1 arrest, energy production requires autophagic degradation of cytosolic components including organelles such as mitochondria (Kang et al., 2007). Sustaining a prolonged search for food during L1 arrest results in the manifestation of aging-like phenotypes including the accumulation of protein aggregates and reactive oxygen species, as well as the fragmentation of the mitochondrial network and depletion of mtDNAs (Hibshman et al., 2018; Roux et al., 2016). Once food is encountered, the aging-like deterioration is resolved and growth resumes (Roux et al., 2016), which is accompanied by increased transcription and translation of components required for mitochondrial biogenesis (Baugh et al., 2009; Stadler and Fire, 2013; Wang and Kim, 2003).

Nuclear chromosome replication is regulated by numerous proteins and checkpoints to ensure that genome duplication is coordinated with the cell cycle (Sclafani and Holzen, 2007; Vermeulen et al., 2003). Akin to nuclear genome replication, mtDNA replication is often followed by mitochondrial fission presumably to ensure that each daughter mitochondrion inherits at least one mtDNA (Lewis et al., 2016). However, mtDNA replication is not coordinated with the cell cycle. While the majority of the proteins required for mtDNA replication have been identified, the upstream events that stimulate mtDNA replication remain unclear. Prolonged L1 arrest and the subsequent recovery offers a physiologically relevant as well as genetically tractable model to study the signaling pathways regulating bulk initiation of mtDNA replication and recovery of the mitochondrial network. Here, we examine the role of the UPR^mt^ and ATFS-1 in coordinating mtDNA content recovery or expansion following prolonged starvation.

## RESULTS

### ATFS-1 is required for development following prolonged starvation at L1 stage

*atfs-1(null)* worms, which lack the entire open reading frame have reduced mitochondrial respiration, function and have shortened adult lifespan (Deng et al., 2019; Shpilka et al., 2021) (Figure 1B). To survive prolonged starvation, mitochondrial function and integrity is required as loss-of-function mutations in the OXPHOS complex III component *isp-1* or in the ubiquinone synthesis component *clk-1* are short-lived during L1 arrest (Lee and Ashrafi, 2008). Thus, we hypothesized that ATFS-1 is also required for survival during starvation. Surprisingly, *atfs-1(null)* worms survived as long as wildtype worms during L1 arrest (Figure 1C). Together, these data indicate that despite the reliance on oxidative phosphorylation, the UPR^mt^ is not involved in mitochondrial maintenance or survival during L1 arrest.

We next examined the growth of wildtype and *atfs-1(null)* worms hatched in the presence of food (referred to as “control”) compared to development upon exposure to food following 5 days of L1 arrest (“referred to as L1 arrest-fed”) (Figure 1D). Strikingly, the development of *atfs-1(null)* worms was significantly impaired following L1 arrest (Figures 1E, 1F), while wildtype worms developed similarly to worms hatched on food (Figures 1E, F). When raised in the presence of food, *atfs-1(null)* worms produce progeny, although fecundity is reduced relative to wildtype worms. However, following starvation *atfs-1(null)* worms did not lay eggs or produce any viable progeny (Figures 1G, Figure S1A). Additionally, *atfs-1(null)* worms appeared more transparent and were smaller compared to wildtype worms of the same stage, suggestive of abnormal development (Figure S1B).

As *atfs-1(null)* worms were sterile following L1 arrest, we examined the germline using transgenic germline reporter strains. The GLH-1::GFP fusion protein marks P-granules which are RNA-enriched compartments specific to germ cells (Seydoux 2008). The germlines of *atfs-1(null)* and wildtype worms hatched in the presence of food expressed GLH-1:GFP throughout the germline (Figures 1H). However, the *atfs-1(null)* worms that were fed following L1 arrest had only a cluster of germ cells expressing GLH-1::GFP that failed to expand (Figure 1H). Further, we used a reporter strain expressing PIE-1::GFP to examine germline proliferation. Germ cells express PIE-1 to prevent the adoption of somatic cell fates by repressing transcription (Mello et al., 1996; Reese et al., 2000). Interestingly, following L1 arrest the germline in *atfs-1(null)* worms did not proliferate to an extent where PIE-1::GFP was expressed (Figure 1I). Combined, these data indicate that the UPR^mt^ is required to resume growth and establish a functional germline following prolonged L1 arrest.

### UPR^mt^ promotes mtDNA content recovery following L1 starvation

As ATFS-1 promotes mitochondrial maintenance during growth, we examined mitochondrial network morphology and function using the dye TMRE which accumulates within functional mitochondria. At the L4 stage, TMRE staining was comparable throughout the soma in wildtype worms fed upon hatching or following 5 days of L1 arrest (Figures 2A, 2B). As expected, *atfs-1(null)* worms accumulated less TMRE than wildtype worms when hatched in the presence food (Shpilka et al., 2021). However, following 5 days of L1 arrest, TMRE staining was nearly undetectable in *atfs-1(null)* worms compared to worms hatched on food of similar age indicating that these worms were unable to establish a functional mitochondrial network (Figures 2A, 2B).

**Figure 2.**
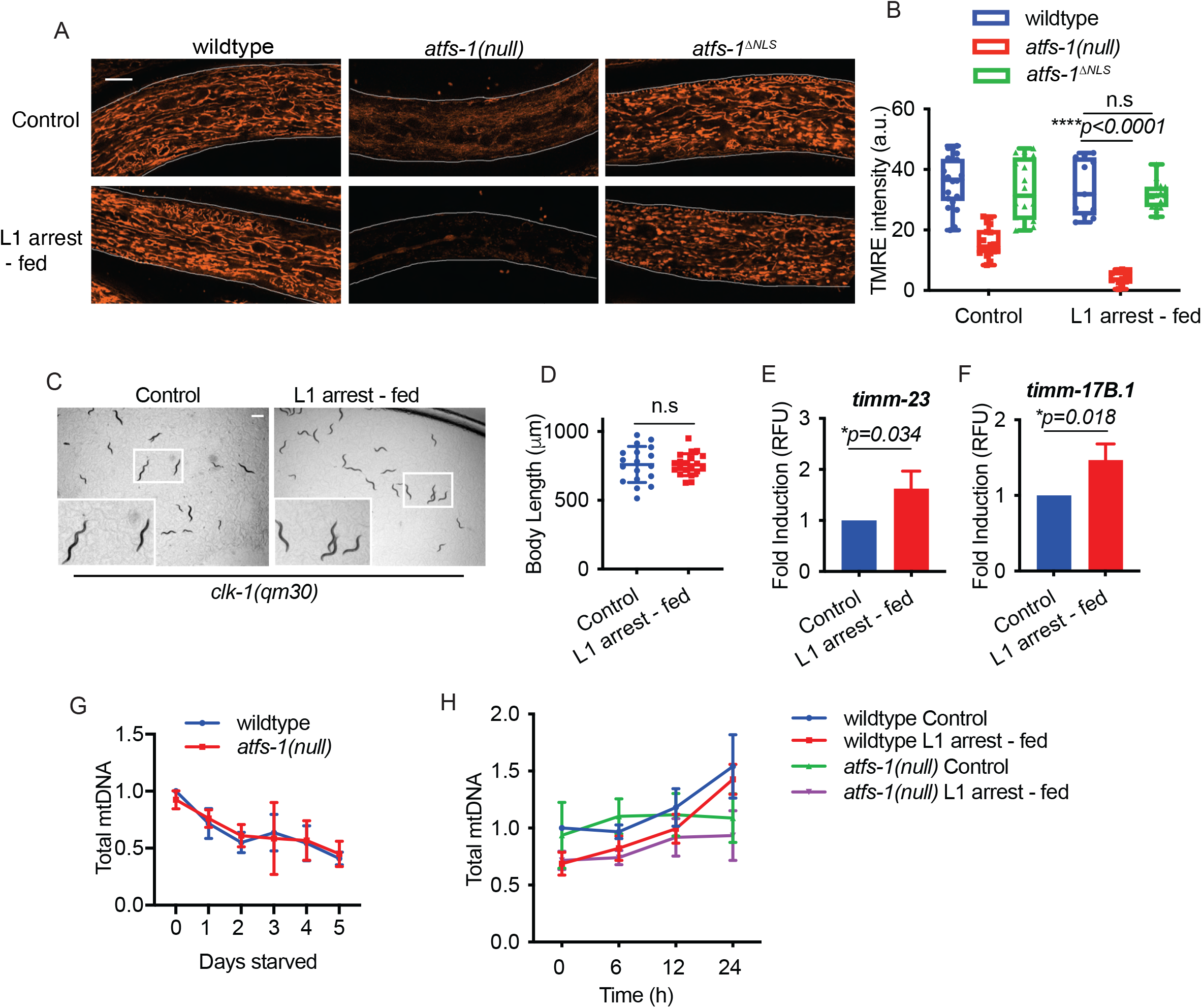
UPR^mt^ promotes mtDNA content recovery following starvation. A. Images comparing TMRE staining of wildtype, *atfs-1(null)* and *atfs-1*Δ^*NLS*^ worms under control or L1 arrest-fed conditions obtained when control worms reached L4 stage (Scale bar, 10 μm, representative images from three biological replicates). B. Quantification of TMRE staining of wildtype, *atfs-1(null)* and *atfs-1*Δ^*NLS*^ worms at the L4 stage in control or L1 arrest-fed conditions (*n = 15 to 20* worms for each strain and condition □±□SD, *\*\*\*\*p < 0*.*0001*, Student’ s *t*-test, a.u., arbitrary units). C. Images comparing development of *clk-1(qm30)* worms in control and L1 arrest-fed conditions on day 1 of adulthood (Scale bar, 0.5 mm). D. Quantification of body lengths of *clk-1(qm30)* worms in control and L1 arrest-fed conditions measured on day 1 of adulthood (*n = 3* ± SD, Student’ s t-test). E. Transcript level of *timm-23* as measured by qRT-PCR at the L2 stage (*n = 3* ± SD, *\*p = 0*.*034*, Student’ s t-test). F. Transcript level of *timm-17B*.*1* as measured by qRT-PCR at the L2 stage (*n = 3* ± SD, *\*p = 0*.*018*, Student’ s t-test). G. mtDNA quantification of wildtype and *atfs-1(null)* starved L1s over a period of 5 days as determined by qPCR. (*n = 3* ± SD, one-way ANOVA, wildtype worms - *\*\*\*p = 0*.*0007*, Post-hoc Dunett’ s test shows significant difference between 0 days starved worms and 1 to 5 days starved worms, *atfs-1(null)* worms – *\*p = 0*.*0191*, Post-hoc Dunett’ s test shows significant difference between 0 days starved worms and 3 to 5 days starved worms). H. Time course of mtDNA content over 24 hours as determined by qPCR comparing wildtype and *atfs-1(null)* worms in control and L1 arrest-fed conditions (*n = 6* for wildtype and *n = 3* for *atfs-1(null)* ± SD, *\*\*p = 0*.*0033* comparing L1 arrest-fed wildtype and *atfs-1(null)* mtDNA content following 24 hours of feeding).

As TMRE staining was undetectable in *atfs-1(null)* worms following starvation, we hypothesized that the impaired recovery from starvation is due to accumulating mitochondrial damage caused by OXPHOS dysfunction. However, we found that *clk-1(qm30)* mutant worms that have impaired respiration, developed at the same rate regardless of whether they were hatched in the presence of food or underwent L1 arrest prior to feeding (Figures 2C, 2D). This data suggests that the developmental defects of *atfs-1(null*) worms are not driven solely by underlying OXPHOS dysfunction but rather by lack of a UPR^mt^ mediated process.

The best studied function of ATFS-1 and the UPR^mt^ is the transcriptional regulation of nuclear genes that promote mitochondrial proteostasis and mitochondria biogenesis (Baker et al., 2012; Shpilka et al., 2021; Yoneda et al., 2004). Consistent with UPR^mt^ activation during recovery from starvation, mRNAs encoding the inner mitochondrial membrane protein import components *timm-23* and *timm-17B*.*1* were found to be increased relative to control worms (Figures 2E, 2F). As a previous ChIP-sequencing experiment demonstrated that ATFS-1 binds to the *timm-23* promoter during mitochondrial stress and thus, likely directly regulates its transcription following L1 arrest (Nargund et al., 2015), these combined findings indicate that ATFS-1-dependent transcription is active during recovery from prolonged L1 arrest.

In addition to reduced transcription of nuclear-encoded genes involved in mitochondrial biogenesis, *atfs-1(null)* worms also harbor fewer mtDNAs than wildtype worms (Shpilka et al., 2021). Thus, we examined the mtDNA content of *atfs-1(null)* worms during L1 arrest and following food introduction. A previous study found that mtDNAs were depleted during L1 arrest by autophagy over time (Hibshman et al., 2018). As expected, mtDNA content was reduced in wildtype worms over the period of 5 days of L1 arrest (Figures 2G). *atfs-1(null)* worms hatched with similar quantities of mtDNA as wildtype worms that also reduced during L1 arrest to ∼50% by day 5 (Figure 2G). Interestingly, while mtDNA content increased in wildtype worms upon food introduction, mtDNA content remained low in *atfs-1(null)* worms (Figure 2H). Combined, these data suggest that once L1 worms begin to feed, ATFS-1 is required to increase, or recover, the mtDNA content that was degraded during L1 arrest (Figure 2G, 2H). These findings emphasize the challenges in mediating the metabolic transition that must occur when the worm goes from a catabolic state to fuel the search for food to an anabolic state required for worm growth and development once feeding begins.

### Mitophagy does not impair mtDNA accumulation in *atfs-1(null)* worms following L1 arrest

Given the reduced mtDNA accumulation in *atfs-1(null)* worms following L1 arrest, we hypothesized that this may be either due to impaired mtDNA replication or increased mitophagy-mediated clearance of mitochondria as a result of accumulating mitochondrial damage in the absence of the UPR^mt^. To determine the impact of mitophagy, we examined recovery from L1 arrest in *atfs-1(null);pdr-1(tm598)* worms that contains a loss-of-function mutation in the ubiquitin ligase Parkin, which is required for mitophagy in *C. elegans* (Lin et al., 2016; Narendra et al., 2008). While *atfs-1(null);pdr-1(tm598)* worms developed slower than either single mutant, they are able to develop to adulthood when hatched in the presence of food (Figure S2A, S2B). However, similar to *atfs-1(null)* worms, the recovery of mtDNA content following L1 arrest was impaired in *atfs-1(null);pdr-1(tm598)* worms and they also developed slowly (Figure S2A, S2B, S2C, S2D). These findings suggest the impaired recovery of mtDNA content in *atfs-1(null)* worms is not due to mitophagy.

### ATFS-1-dependent nuclear transcription is not required for mtDNA recovery

Our findings suggest that recovery of mtDNA copy number following starvation could be due to ATFS-1 1) directly promoting mtDNA replication and/or 2) regulating nuclear transcription that promotes mitochondria function. To examine the role of ATFS-1-dependent nuclear transcription, we compared wildtype, *atfs-1(null)* and mutant worms in which the nuclear localization sequence (NLS) within ATFS-1 was mutated (Shpilka et al., 2021; Yang et al., 2022). Consistent with impaired nuclear activity of *atfs-1*^Δ*NLS*^ worms, previous studies showed that the increase in *atfs-1-*dependent transcription of the nuclear gene *hsp-6* was reduced in *atfs-1*^Δ*NLS*^ worms raised on mitochondrial stress (Shpilka et al., 2021; Yang et al., 2022). Importantly, ATFS-1^Δ*NLS*^ is imported into mitochondria and processed similarly to wildtype ATFS-(Yang et al., 2022). *atfs-1*^Δ*NLS*^ worms had similar lifespans as wildtype worms while the lifespan of *atfs-1(null)* worms was considerably reduced (Figure 1B). Further, we generated *clk-1(qm30);atfs-1*^Δ*NLS*^ worms to examine the impact of reduced *atfs-1*-dependent nuclear transcription during mitochondrial dysfunction. We found that *clk-1(qm30);atfs-1*^Δ*NLS*^ worms developed considerably slower and produced fewer progeny than *clk-1(qm30)* worms indicating that the nuclear activity of ATFS-1 is required for growth during OXPHOS perturbation (Figures 3A, 3B).

**Figure 3.**
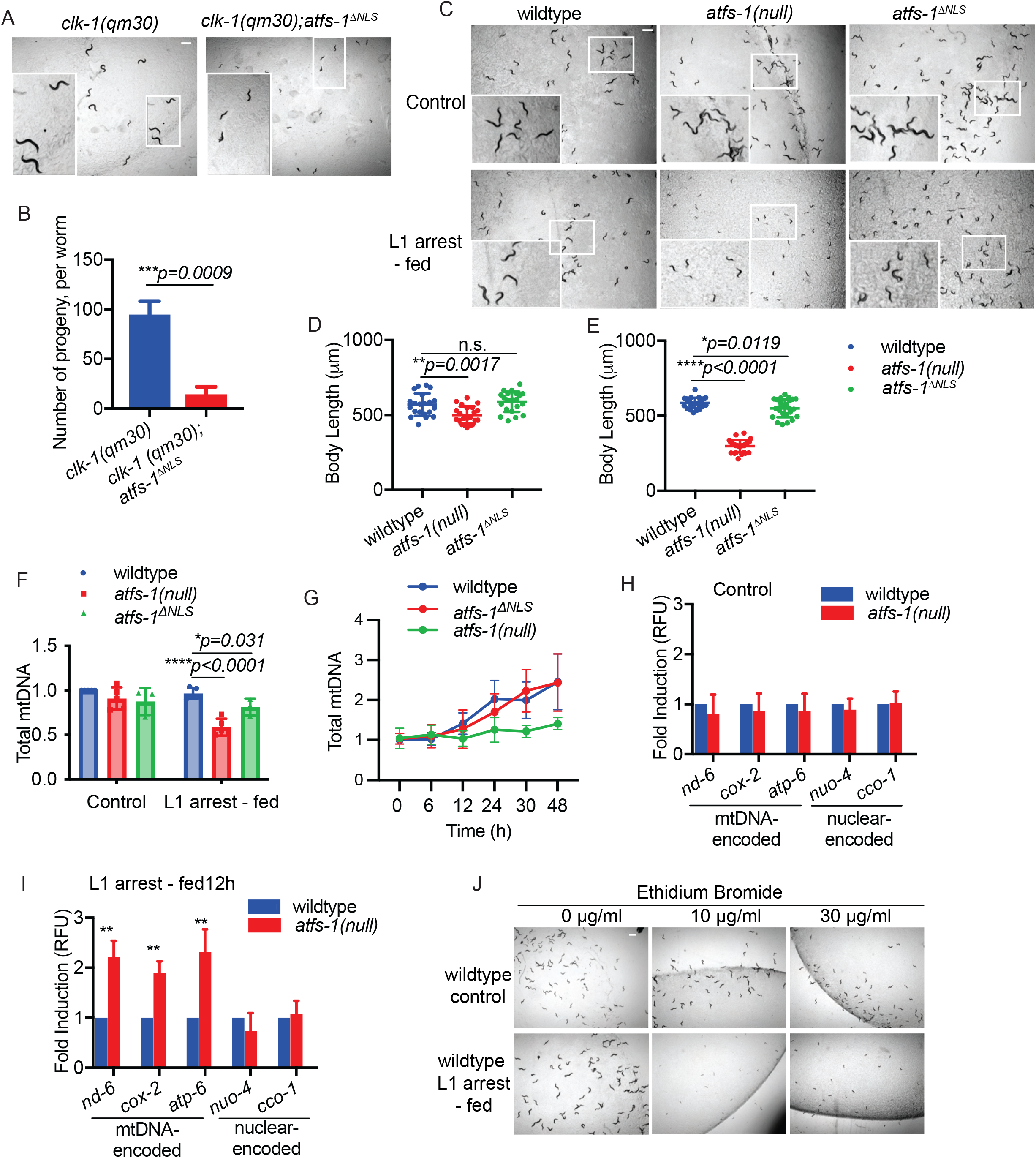
The increase in mtDNA content following L1 arrest does not require *atfs-1*-dependent nuclear transcription. A. Images comparing the size of *clk-1(qm30)* and *clk-1(qm30);atfs-1*^Δ*NLS*^ worms hatched on food obtained when the *clk-1(qm30)* reached day 1 of adulthood (Scale bar, 0.5 mm). B. Brood size comparison of *clk-1* or *clk-1(qm30);atfs-1*^Δ*NLS*^ worms in control or L1 arrest-fed conditions (*n = 3* ± SD, *\*\*\*p = 0*.*0009*, Student’ s t-test). C. Images of wildtype, *atfs-1(null)* and *atfs-1*^Δ*NLS*^ worms in control and L1 arrest-fed conditions obtained when the control condition worms reached the L4 stage (Scale bar, 0.5 mm). D. Quantification of body lengths of wildtype, *atfs-1(null)* and *atfs-1*^Δ*NLS*^ worms in control condition when the worms reached L4 stage (*n = 3* ± SD, *\*\*p = 0*.*0017*, Student’ s t-test). E. Quantification of body lengths of wildtype, *atfs-1(null)* and *atfs-1*^Δ*NLS*^ worms in L1 arrest-fed condition when the control worms reached the L4 stage (*n = 3* ± SD, *\*p = 0*.*0119* comparing wildtype and *atfs-1*^Δ*NLS*^ worms and *\*\*\*\*p < 0*.*0001* comparing wildtype and *atfs-1(null)* worms, Student’ s t-test). F. mtDNA quantification of wildtype, *atfs-1(null)* and *atfs-1*^Δ*NLS*^ worms when the control condition worms reached the L4 stage (*n = 3 to 5* ± SD, *\*\*\*\*p < 0*.*0001* comparing wildtype and *atfs-1(null)* worms and *\*p = 0*.*031* comparing wildtype and *atfs-1*^Δ*NLS*^ L1 arrest - fed worms, Student’ s t-test). G. Time course of mtDNA quantification by qPCR comparing wildtype, *atfs-1(null)* and *atfs-1*^Δ*NLS*^ worms fed for 48 hours following L1 arrest (*n = 4* ± SD for wildtype *\*p = 0*.*0264* comparing wildtype and *atfs-1(null)* worm mtDNA at 48 hours, *\*p = 0*.*0305* comparing wildtype and *atfs-1*^Δ*NLS*^ worm mtDNA at 48 hours, Student’ s t-test). H. Transcript levels of mtDNA- and nuclear DNA-encoded OXPHOS genes in wildtype and *atfs-1(null)* control worms 24 hours after hatching on food (*n = 3* ± SD, Student’ s t-test). I. Transcript levels of mtDNA and nuclear DNA-encoded genes in wildtype and *atfs-1(null)* L1 arrest-fed worms at 12 hours following the introduction of food (*n = 3* ± SD, *\*\*p* ≤ *0*.*01*, Student’ s t-test). J. Images of wildtype control and L1 arrest-fed worms treated with 0, 10 and 30 μg/ml of ethidium bromide when the control condition worms reached the L4 stage (Scale bar, 0.5 mm).

We next examined the role of ATFS-1 nuclear activity during development following prolonged L1 arrest. Strikingly, *atfs-1*^Δ*NLS*^ worms developed much faster than *atfs-1(null)* worms (Figures 3C, 3D, 3E). Interestingly, upon feeding following prolonged L1 arrest, mtDNA content recovered at similar rates in wildtype and *atfs-1*^Δ*NLS*^ worms but the recovery was impaired in *atfs-1(null)* worms (Figure 3F, 3G). Importantly, unlike *atfs-1(null)* worms, *atfs-1*^Δ*NLS*^ worms established a functional mitochondrial network upon feeding as determined by TMRE staining (Figure 2A). However, the mitochondrial network was more fragmented in *atfs-1*^Δ*NLS*^ worms relative to wildtype worms (Figure S3A) indicating that *atfs-1-*dependent transcription is required to maintain optimal mitochondrial function following prolonged starvation.

To further examine the role of mitochondrial-localized ATFS-1 in recovery from L1 arrest, we evaluated the mitochondrial network in *atfs-1*^*R/R*^ worms in which the strength of the ATFS-1 mitochondrial targeting sequence was increased by substituting the threonine and aspartic acid residues at positions 10 and 24 with arginine residues (Shpilka et al., 2021). Consistent with increased import of ATFS-1^R/R^ into mitochondria, *atfs-1*^*R/R*^ worms have an impaired nuclear response during mitochondrial stress and a fragmented mitochondrial network (Shpilka et al., 2021). Similar to *atfs-1*^Δ*NLS*^ worms, L1 arrested *atfs-1*^*R/R*^ worms developed to adulthood upon feeding (Figure S3B). Importantly, the mtDNA content in *atfs-1*^*R/R*^ worms was also higher than in *atfs-1(null)* worms exiting prolonged L1 arrest (Figure S3C). Combined, these findings indicate that mitochondrial-localized ATFS-1 is required to resume growth and recover mtDNA content following prolonged L1 arrest.

We previously found that ATFS-1 binds directly to mtDNA during mitochondrial dysfunction and inhibition of *atfs-1* during mitochondrial stress resulted in an aberrant increase of mtDNA-encoded OXPHOS transcripts (Nargund et al., 2015). We hypothesized that in addition to promoting mtDNA recovery, mitochondrial-localized ATFS-1 might also limit mtDNA-encoded transcript accumulation to promote development following starvation. Interestingly, we found that several mtDNA-encoded mRNAs were increased in *atfs-1(null)* worms, but not nuclear genome-encoded oxidative phosphorylation (OXPHOS) genes following the introduction of food (Figures 3H, 3I). These findings suggest that in addition to promoting mtDNA replication, mitochondrial-localized ATFS-1 represses or limits transcription of mtDNA-encoded OXPHOS transcripts following prolonged L1 starvation potentially to facilitate efficient OXPHOS complex assembly (Nargund et al., 2015).

We next sought to determine whether the *atfs-1*-mediated recovery of mtDNA content or the repression of mtDNA-encoded OXPHOS mRNAs is required for the recovery from prolonged L1 arrest. Thus, we exposed L1 arrested wildtype worms to ethidium bromide (EtBr), which intercalates within mtDNA and inhibits replication (Martinus et al., 1996; Yoneda et al., 2004). The exposure to 10µg/ml and 30µg/ml EtBr did not affect development of wildtype worms hatched directly onto food (Figure 3J). Impressively, the same doses severely impaired the development of worms following prolonged L1 arrest (Figure 3J). Combined, these data suggest that the recovery of mtDNA copy number mediated by mitochondrial-localized ATFS-1 is a limiting factor for growth and germline development following prolonged L1 arrest.

### *daf-2* inhibition promotes *atfs-1*-dependent mtDNA replication

Lastly, we sought to gain insight into the upstream events regulating *atfs-1*-dependent mtDNA replication that occurs following prolonged L1 arrest. We and others previously found that inhibition of the insulin-like receptor DAF-2 using a hypomorphic mutant (*daf-2(e1370)*), impairs *atfs-1*-dependent induction of *hsp-6*_*pr*_*::gfp* during mitochondrial stress (Figure 4A) suggesting that DAF-2 activity promotes nuclear accumulation of ATFS-1 (Gatsi et al., 2014; Shpilka et al., 2021). Interestingly, *daf-2(e1370)*, hatched in the presence of food harbored nearly 1.5-fold more mtDNAs than wildtype worms at the L4 stage suggesting that modest inhibition of DAF-2 promotes mtDNA replication (Haroon et al., 2018) (Figure 4B). We hypothesized that the increased mtDNA content of *daf-2(e1370)* mutant was dependent on ATFS-1. Interestingly, we found that the increase in mtDNAs was abrogated in *daf-2(e1370);atfs-1(null)* mutants (Figure 4B). Further, *daf-2(e1370);atfs-1*^Δ*NLS*^ worms harbored similar amounts of mtDNAs as *daf-2(e1370)* worms (Figure 4C) suggesting mitochondrial-localized ATFS-1 promotes mtDNA replication when *daf-2* is inhibited. Consistent with ATFS-1 localizing to the mitochondria, a chromatin precipitation (ChIP) assay using ATFS-1-specific antibodies (Nargund et al., 2015) indicated that ATFS-1 interacts with 2-fold more mtDNAs upon DAF-2 inhibition (Figure 4D). Together these data suggest that reduced DAF-2 activity diminishes nuclear function of ATFS-1 which allows mitochondrial-localized ATFS-1 to promote an increase in mtDNA content.

**Figure 4.**
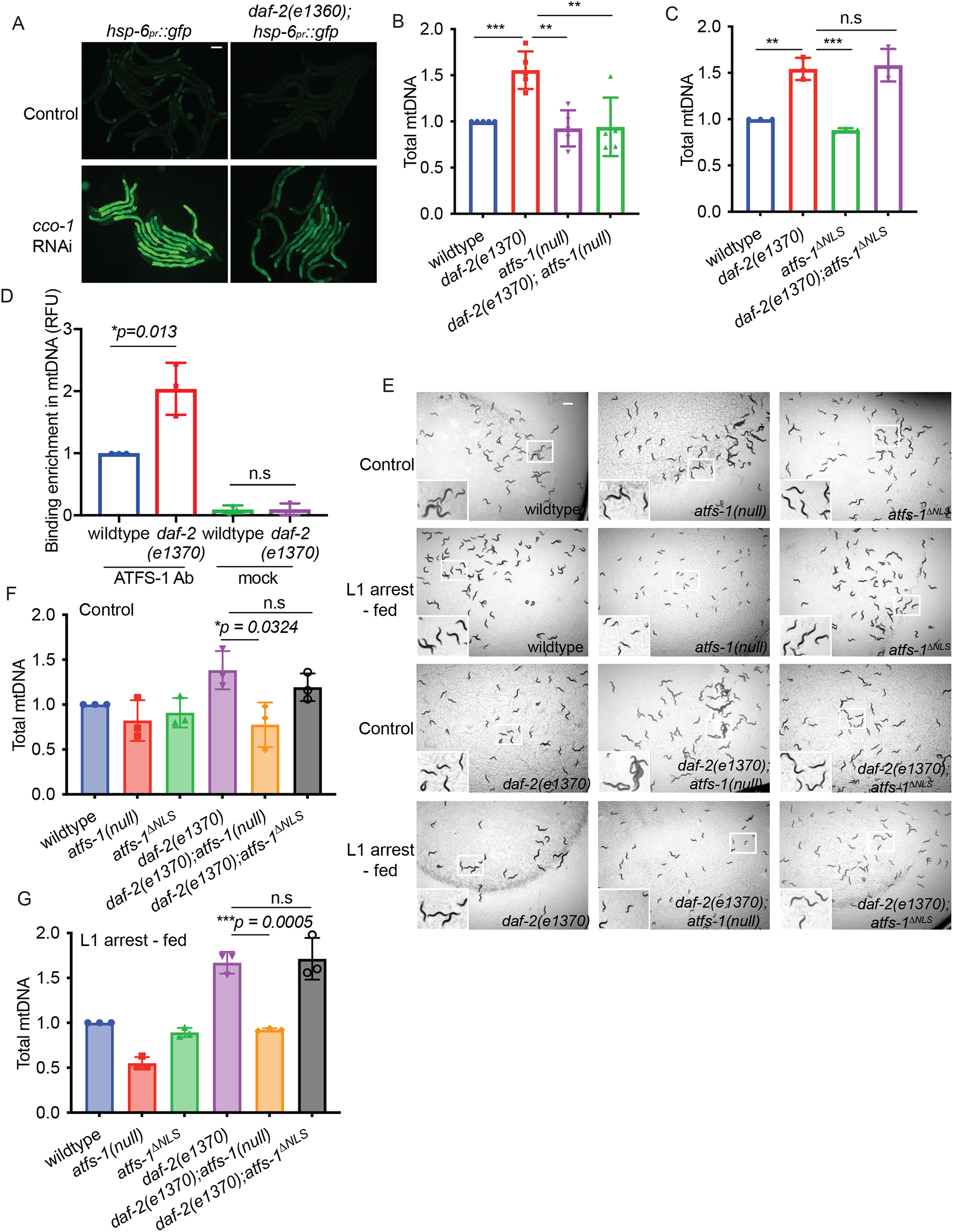
*daf-2* inhibition promotes *atfs-1*-dependent mtDNA replication. A. Fluorescence images of *hsp-6*_*pr*_*::gfp* and *daf-2(e1370);hsp-6*_*pr*_*::gfp* worms raised on control or *cco-1*(RNAi) at the L4 stage (Scale bar, 0.1 mm). B. mtDNA quantification of wildtype, *atfs-1(null), daf-2(e1370)* and *daf-2(e1370);atfs-1(null)* worms determined at the L4 stage (*n = 5* ± SD, \**\*\*p* = *0*.*0003* comparing wildtype and *daf-2(e1370)* worms, *\*\*p* = *0*.*0065* comparing *daf-2(e1370)* and *daf-2(e1370);atfs-1(null)* worms, *\*\*p=0*.*0011* comparing *daf-2(e1370)* and *atfs-1(null)* worms, Student’ s t-test). C. mtDNA quantification of wildtype, *atfs-1*^Δ*NLS*^, *daf-2(e1370)* and *daf-2(e1370);atfs-1*^Δ*NLS*^ worms determined at the L4 stage (*n = 3* ± SD, *\*\*p* = *0*.*0015* comparing wildtype and *daf-2(e1370)* worms, *\*\*\*p = 0*.*0007* comparing *daf-2(e1370)* and *atfs-1*^Δ*NLS*^ worms, Student’ s t-test). D. mtDNA quantification following ATFS-1- or control-ChIP in wildtype and *daf-2(e1370)* worms at the L4 stage (*n = 3* ± SD, *\*p = 0*.*013*, Student’ s t-test). E. Images of wildtype, *atfs-1(null), atfs-1*^Δ*NLS*^, *daf-2(e1370), daf-2(e1370);atfs-1(null) and daf-2(e1370);atfs-1*^Δ*NLS*^ worms in control and L1 arrest-fed conditions, obtained when control worms reached L4 stage (Scale bar, 0.5 mm). F. mtDNA quantification of wildtype, *atfs-1(null), atfs-1*^Δ*NLS*^, *daf-2(e1370), daf-2(e1370);atfs-1(null) and daf-2(e1370);atfs-1*^Δ*NLS*^ worms in control condition as determined by qPCR at the L4 stage (*n = 3* ± SD, *\*p = 0*.*0324* comparing *daf-2(e1370)* and *daf-2(e1370;atfs-1(null)* worms, Student’ s t-test). G. mtDNA quantification of wildtype, *atfs-1(null), atfs-1*^Δ*NLS*^, *daf-2(e1370), daf-2(e1370);atfs-1(null)* and *daf-2(e1370);atfs-1*^Δ*NLS*^ worms in L1-arrest condition as determined by qPCR when control worms reached the L4 stage (*n = 3* ± SD, *\*\*\*p = 0*.*0005* comparing *daf-2(e1370)* and *daf-2(e1370;atfs-1(null)* worms, Student’ s t-test).

Next, we examined the effect of L1 arrest on the mtDNA content of *daf-2(e1370)* mutants. While *daf-2(e1370)* worms hatched with increased mtDNA content relative to wildtype worms, mtDNA content declined at similar rates in both strains during L1 arrest (Figure S4A). We hypothesized that the increased mtDNA content of *daf-2(e1370)* throughout L1 arrest can be advantageous for growth or recovery following prolonged L1 arrest. As expected, growth was impaired in wildtype worms exposed to EtBr while *daf-2(e1370)* worms developed to adulthood following prolonged L1 arrest (Figure S4B) further suggesting that the lack of mtDNA is indeed a rate limiting factor for growth following starvation.

We next examined the role of ATFS-1 in the recovery of *daf-2(e1370)* worms following prolonged starvation. Interestingly, *daf-2(e1370);atfs-1(null)* worms did not develop into mature adults (Figure 4E). However, *daf-2(e1370);atfs-1*^Δ*NLS*^ worms developed into fertile adults similarly to wildtype worms (Figure 4E). Consistent with these results, *daf-2(e1370);atfs-1(null)* worms harbor fewer mtDNAs relative to *daf-2(e1370)* and *daf-2(e1370);atfs-1*^Δ*NLS*^ worms following starvation suggesting that ATFS-1 functions downstream of DAF-2 inhibition to increase the mtDNA content (Figures 4F, 4G) required for development following prolonged L1 arrest.

The FOXO-like transcription factor DAF-16 is the canonical downstream effector activated upon DAF-2 inhibition. DAF-16 promotes L1 survival by regulating transcription of numerous genes including the cyclin-dependent kinase inhibitor and by altering metabolic flux to promote trehalose synthesis during L1 arrest (Baugh and Sternberg, 2006; Hibshman et al., 2017). Moreover, *daf-16* loss-of-function mutant worms are unable to develop following prolonged L1 arrest (Marta Sanz, 2019). Interestingly, *daf-2(e1370);daf-16(mu86)* worms had fewer mtDNAs than *daf-2(e1370)* worms consistent with DAF-16 also promoting mtDNA accumulation (Figure S4C). Thus, we sought to understand the relationship between *daf-16 and atfs-1* using a partial deletion mutant *daf-16(mu86)*. Importantly, *daf-2(e1370);daf-16(mu86);atfs-1(null)* worms had a further reduction in mtDNAs compared to either single deletion suggesting that ATFS-1 and DAF-16 function in independent pathways downstream of DAF-2 to regulate mtDNA content (Figure S4D). As *daf-*16-mutants are unable to maintain and survive prolonged L1 arrest (Baugh and Sternberg, 2006; Hibshman et al., 2017), the role of DAF-16 in increasing mtDNA content upon feeding remains unclear. Based on these data, we concluded that a reduction in DAF-2 activity leads to increased mtDNA content dependent on mitochondrial ATFS-1 that was sufficient to promote growth and mitochondrial recovery after prolonged starvation.

## DISCUSSION

*C. elegans* mature from egg to reproductive adult within three days. However during development, worms can enter a diapause-like state at different developmental stages to endure prolonged stresses including starvation (Baugh and Hu, 2020). Those worms that hatch in the absence of food remain developmentally arrested at the L1 stage and can survive for over thirty days (Roux et al. (2016), Figure 1C). To sustain the search for food, cytosolic components including mitochondria are degraded by autophagy which is used to generate ATP via oxidative phosphorylation (Hibshman et al., 2018; Kang et al., 2007). Once food is encountered, a major challenge to the resumption of development is repairing the cellular and tissue damage incurred in the absence of food (Roux et al., 2016).

Here, we find that following five days of L1 arrest ATFS-1 and the UPR^mt^ are required for development and to establish a functional germline. A previous study found a strong correlation between those mutations that extend L1 survival and those mutations that extend adult lifespan (Muñoz and Riddle, 2003). In contrast, we found that while *atfs-1(null)* worms are relatively short-lived when hatched in presence of food, however the absence of ATFS-1 does not impact L1 survival. Moreover, ATFS-1 does not appear to have any role in the mtDNA accumulation that occurs during embryogenesis through L1 stage. During prolonged L1 arrest, mtDNAs and presumably whole mitochondria are degraded via autophagy (Hibshman et al., 2018). Our findings indicate that mtDNA depletion occurs independent of ATFS-1 during L1 arrest. However, upon the introduction of food, ATFS-1 is required to increase mtDNA content to a level comparable to that of newly hatched worms. To our surprise, the NLS within ATFS-1 required for *atfs-1*-dependent nuclear transcription was not required to recover mtDNA content or establish a functional mitochondrial network or functional germline. However, it should be noted that *atfs-1*^Δ*NLS*^ worms have perturbed mitochondrial morphology at the L4 stage, consistent with ATFS-1 coordinating mitochondrial network expansion with the increased levels of protein synthesis and growth that occurs upon feeding (Shpilka et al., 2021).

Lastly, our findings suggest that the activity of mitochondrial-localized ATFS-1 is required to increase in mtDNA content upon inhibition of the insulin receptor-like protein DAF-2. We previously found that *daf-2*-inhibition impairs *atfs-1-*dependent transcription, likely due to impairing S6 kinase and reducing general rates of protein synthesis which results in the majority of ATFS-1 trafficking to mitochondria (Shpilka et al., 2021). Here, we find that a reduction in DAF-2 function causes an increase in mtDNA content which requires *atfs-1*, but not *atfs-1-* dependent nuclear transcription, suggesting that a reduction in DAF-2 activity promotes mtDNA replication via mitochondrial-localized ATFS-1, but limits nuclear transcription. We propose that this “toggle switch”-like activity of DAF-2 ensures the synthesis of mtDNAs occurs prior to activation of the transcription program required for the expansion of the mitochondrial network driven by TORC1-dependent protein synthesis during worm development. It is interesting to note that these findings are consistent with our previous work indicating that mitochondrial-localized ATFS-1 promotes mtDNA replication in dysfunctional mitochondria, which can contribute to a replicative advantage of deleterious mtDNAs in a disease model (Lin et al., 2016; Yang et al., 2022). Our findings that ATFS-1 is required to increase mtDNA replication and recover mtDNAs degraded during starvation suggest a reason why such a pathway may have evolved, as recovery of mtDNA content following L1 arrest is essential for development and *C. elegans* reproduction.

Previous studies demonstrated that starvation leads to a reduction of *daf-2*-mediated signaling as DAF-16 activity is required for starvation survival as well as maintaining germline quiescence (Baugh and Sternberg, 2006; Hibshman et al., 2017). Furthermore, upon feeding, *daf-2* transcription is reduced, and DAF-16 activity is necessary to promote growth and development (Chen and Baugh, 2014; Kaplan et al., 2019; Olmedo et al., 2020) suggestive of a more nuanced *daf-2* regulation. We propose a model that the prolonged lack of food during L1 arrest leads to modest reduction of *daf-2* signaling along with the gradual decline in mitochondrial activity while sustaining foraging, depletion of mtDNA, mitochondrial fragmentation and reduced respiration. Upon feeding, the dysfunctional mitochondrial network combined with reduced DAF-2 signaling initiates mtDNA replication via ATFS-1 that precedes or coincides with the transcription of nuclear genes required for mitochondrial biogenesis.

## ACKNOWLEGEMENTS

We thank the Caenorhabditis Genetics Center for providing *C. elegans* strains funded by NIH Office of Research 362 Infrastructure Programs (P40 OD010440).

## Author Contributions

N.U.N, T.S and C.M.H. designed the experiments. N.U.N, Y.D, and Q.Y., generated the worm strains and performed research. N.U.N and C.M.H. wrote the manuscript.

## STAR Methods

### Key resources Table

| REAGENT or RESOURCE | SOURCE | IDENTIFIER |
| --- | --- | --- |
| **Experimental models: Organisms/strains** |  |  |
| *C. elegans*: MQ130 (*clk-1(qm30)*) | Caenorhabditis Genetics Center (CGC) | N/A |
| *C. elegans*: CB1370 (*daf-2(e1370)*) | Caenorhabditis Genetics Center (CGC) | N/A |
| *C. elegans*: CB1038 (*daf-16(mu86)*) | Caenorhabditis Genetics Center (CGC) | N/A |
| *atfs-1*<sup>ΔNLS</sup> | Shpilka et al., 2021, Yang et al., 2022 | N/A |
| *atfs-1*<sup>R/R</sup> | Shpilka et al., 2021 | N/A |
| *pdr-1(tm598)* | National BioResource Project (Tokyo, Japan) | N/A |
| *atfs-1(null)* | Deng et al., 2019 | N/A |
| PIE-1::GFP | Laboratory of Dr. Craig C. Mello | N/A |
| GLH-1::GFP | Laboratory of Dr. Craig C. Mello | N/A |
| **Oligonucleotides** |  |  |
| Full list of qRT-PCR primers is presented in Table S2 | This paper | N/A |
| **Chemicals** |  |  |
| Ethidium Bromide | Invitrogen | 15585011 |
| TMRE | Thermo Fisher | T669 |
| **Software and algorithms** |  |  |
| Fiji Image J | Image J | RRID:SCR_002285 |
| Prism | Graphpad | RRID:SCR_002798 |
| OASIS2 | Han et al., 2016 | https://sbi.postech.ac.kr/oasis2/surv/ |

### Lead contact for resource sharing

Further information and requests for reagents or strains must be directed to and will be fulfilled by the lead contact, Cole Haynes.

### Materials availability

This study did not generate new unique materials.

### Method details

***C. elegans* strains** *C. elegans* strains were fed and cultured on HT115 *Escherichia Coli* strain at 20°C unless otherwise specified. The reporter strain *hsp-6*_*pr*_*::gfp* was used for visualizing UPR^mt^ activation (Yoneda et al., 2004). N2 (wildtype), *clk-1(qm30), daf-2(e1370), daf-16(mu86*) strains were obtained from the CGC. PIE::GFP and GLH-1::GFP were gifts from Dr. Craig Mello. *pdr-1(tm598)* was obtained from the National BioResource project (Tokyo, Japan). *atfs-1(null), atfs-1*^Δ*NLS*^, and the *atfs-1*^*R/R*^ strains were generated using CRISPR-Cas9 as described (Deng et al., 2019; Shpilka et al., 2021; Yang et al., 2022).

### Starvation plates and recovery from prolonged L1 arrest

For L1 arrest, worms were synchronized by bleaching and transferred to NGM plates that lacked cholesterol and peptone for 5 days (Stiernagle, 2006). Following L1 arrest, the worms were transferred to NGM plates seeded with HT115 bacteria (L1 arrest-fed condition). Control condition worms were synchronized by bleaching and allowed to hatch directly on NGM plates seeded with HT115 bacteria. All strains were maintained at 20°C throughout all experiments.

For images of worm development, the L1 arrest-fed worms of each strain were compared to control worms of the same developmental stage.

### Brood size quantification

Three L4 stage nematodes were randomly selected from each condition and transferred to a fresh plate and incubated at 20°C. Once worms reached adulthood, the number of progenies were quantified until the worms stopped laying eggs. Brood size was determined as the average number of progenies per worm on each plate based on the sum of total eggs. Three replicates per strain per condition were used.

### mtDNA quantification

mtDNA was measured using quantitative PCR (qPCR) similar to previously published methods (Lin et al., 2016; Shpilka et al., 2021; Valenci et al., 2015). 30-40 worms were collected per sample in lysis buffer containing 50□mM KCl, 10□mM Tris-HCl pH 8.3, 2.5□mM MgCl_2_, 0.45% NP-40, 0.45% Tween 20, 0.01% gelatin, fresh stock of 200□µg/ml proteinase K) and immediately frozen at -80°C for at least 15 minutes followed by lysis at 65°C for 80 minutes. 2μl of lysate was used per reaction, and they were performed in biological and technical triplicates using iQ™ SYBR® Green Supermix and the Biorad qPCR CFX96™ (Bio-Rad Laboratories). The average C_t_ (threshold cycle) of technical replicate values were obtained for mtDNA and was normalized using the 2^−Δ^^Δ^^C^_t_ method. Primers that specifically amplify mtDNA are listed in Table S2. Primers that amplify a non-coding region near the nuclear-encoded *ges-1* gene were used as a control. Student’ s *t*-test or one-way ANOVA was used to determine the statistical significance where applicable.

For measuring mtDNA recovery in Figure 2H over time course of 24 hours, control worms were synchronized to L1s overnight prior to transferring on bacteria seeded plates and L1 arrest-fed worms were starved for 5 days prior to transferring on bacteria seeded plates.

### RNA isolation and qRT-PCR

Worms were harvested from plates into tubes and washed 3 times using *S. basal* to remove bacteria. 1 ml of TRIzol™ Reagent (Invitrogen) was added to the worm pellet and the tubes were flash frozen using liquid nitrogen. Total RNA was isolated from worm pellets using the TRIzol™ Reagent and chloroform. cDNA was then synthesized from total RNA using the iScript cDNA Synthesis Kit (Bio-Rad). qPCR was performed using iQ™ SYBR® Green Supermix. Primer sequences are listed in Table S2. Relative expression of target genes was normalized to the house keeping gene *act-3*. Fold changes in gene expression were calculated.

For qPCR of L1 worms, liquid cultures of different worm strains were grown for at least two generations with food and then prepped for eggs using bleach solution to obtain synchronized L1s. For L1 arrest-fed condition, eggs were hatched onto starvation plates for 5 days and then fed up to L2s before harvesting for RNA extraction. For control condition, eggs directly hatched on food were grown up to L2s were collected and used for RNA extraction.

### Body length measurements

Body length quantifications were obtained at the L4 stage. However, as the *clk-1(qm30)* and *clk-1(qm30); atfs-1*^Δ*NLS*^ strains developed slowly, body length measurements were obtained on day1 of egg lay. Body length measurements were obtained using ImageJ measuring from nose tip to worm tail as previously described (Morck and Pilon, 2006).

### Microscopy

*C. elegans* were imaged using either a Zeiss AxioCam 506 mono camera mounted on a Zeiss Axio Imager Z2 microscope, a Zeiss AxioCam MRc camera mounted on a Zeiss SteREO Discovery V12 stereoscope or a ZEISS LSM800 confocal microscope. Exposure times were the same for each sample in each experiment.

### L1 starvation survival

L1 survival assay was adapted from protocol previously described (Lee and Ashrafi, 2008). Worms were grown for at least two generations at 20°C in liquid with food and bleached to collect eggs, which hatched overnight on a shaker incubator in S-basal without cholesterol. Synchronized L1s were maintained in 25- or 125-ml flasks at a concentration of ∼1 worm/µl in 10–12 ml of S-basal without cholesterol and without antibiotics. Flasks were shaken at 200 rpm in a shaker at 20°C except when scoring. Viability was assessed by aliquoting 100 to 200 µl of worms to fresh plates and monitoring the number of worms that were alive or dead after 2 days post plating. The worms were scored every other day and the survival percent was calculated based on a ratio of the number of worms survived to total number of plated worms on each scoring day. The graph shows data collected from one biological replicate representative of 3 independent replicates. Summary statistics of 3 independent biological replicates are reported in Table S1.

### Adult Lifespan assay

Animals were raised on seeded NGM plates at 20°C until the L4 stage and ∼100 L4 worms were transferred to fresh bacterial plates (10 worms per plate). Worms were scored every 2-3 days and those that were unresponsive to touch of the worm pick were marked as dead. Worms that bagged or crawled off the plate were censored and not included in the study. The figure shows data collected from a single experiment. Each experiment was repeated in 3 independent biological replicates (Table S3).

### DAPI staining

Animals were grown on control or L1 arrest-fed condition until day 1 adulthood. Then, the worms were washed off the plates and placed in microcentrifuge tube containing 0.01% Tween-PBS. Worms were pelleted by low-speed centrifugation and washed 2-3 times with 0.01% Tween PBS. Next, 1 ml of pre-chilled methanol from -20°C was added to the whole worm pellet and tubes were immediately placed in the freezer at -20°C for exactly 5 minutes. The worms were pellet again and rinsed 2-3 times with 0.1% Tween-PBS. 0.2 µl of DAPI solution was added and the tube was allowed in sit in the dark for 5 minutes and after, rinsed with 0.1% Tween-PBS. Finally, 30 µl of 75% glycerol solution was added to worm pellet and were ready to be imaged.

### TMRE staining

Freshly seeded HT115 bacterial plates were coated with TMRE to make a final concentration of 30 µm in each plate. The worms were either synchronized by bleaching and plated onto TMRE plates either from egg stage (control) or starved in L1 stage for 5 days using starvation plates and then transferred to TMRE plates (L1 arrest-fed). Three independent biological repeats were performed. The graph was generated by quantifying of 15-25 individual worms from a single experimental repeat that is representative of all three biological replicates. TMRE quantification was done as previously described (Shpilka et al., 2021). In brief, the average pixel intensity values were calculated by sampling images of different worms. The average pixel intensity for each animal was calculated using ImageJ (http://rsb.info.nih.gov/ij/). Mean values were compared using Student’ s *t-*test.

### Ethidium Bromide experiments

Control worms were bleached directly onto plates containing 0, 10 or 30 µg/ml ethidium bromide with HT115 bacteria. For starved-fed condition, worms were first bleached onto starvation plates for 5 days and then transferred onto ethidium bromide plates as described above. Plates were imaged when at the L4 stage or when they became developmentally arrested.

### ChIP-mtDNA

ATFS-1 ChIP was performed as previously described (Yang et al., 2022). Synchronized worms were cultured on plates and harvested at the L4 stage. The worms were lysed via Teflon homogenizer in chilled PBS with protease inhibitors (Roche). Cross-linking of DNA and protein was performed by treating worms with 1.85% formaldehyde containing protease inhibitors for 15 minutes. Glycine was added to a final concentration of 125 mM and incubated for 5 minutes at room temperature to quench the formaldehyde. The tubes were cold centrifuged at 3000 rpm for minutes and then, pellets were resuspended twice in cold PBS with protease inhibitors. Samples were treated with lysis buffer containing 0.1% Sarkosyl, protease inhibitors in FA Buffer (50 mM HEPES/KOH pH 7.5, 1mM EDTA, 1% Triton X-100, 0.1% Sodium deoxycholate, 150 mM NaCl). The supernatant was precleaned with pre-blocked ChIP-grade Pierce™ magnetic protein A/G beads (Thermo Scientific) and then incubated with Monoclonal Mouse mAb IgG1 Isotype Control (Cell Signaling Technology, G3A1) rotating overnight at 4°C. The antibody-DNA complex was precipitated with protein A/G magnetic beads or protein A sepharose beads (Invitrogen). After washing, the crosslinks were reversed by incubation at 65°C overnight. The samples were then treated with RNaseA at 37°C for 1.5 hour followed by proteinase K treatment at 55°C for 2 hours. Lastly, the immunoprecipitated and input DNA were purified with ChIP DNA Clean & Concentrator (Zymo Research, D5205) and used as templates for qPCR.

### Statistics

All experiments were performed at least three times and all the represented graphs are based on biologically independent replicates. L1 starvation survival and adult life span are graphs from a single biological replicate representative of three independent biological repeats. The summary statistics from three biological replicates are reported in Table S1 and S3. For lifespan analysis, statistics were performed using OASIS2 software (Han et al., 2016) and values were calculated with the log-rank/Mantel Cox test. p-values were calculated by the two-tailed Student’ s t-test. The statistical tests were chosen based on previous studies with similar methodologies and analysis performed using Graphpad Prism 7. Experiments were not blinded, and randomization was not used. For all figures, the mean[±[SD is represented unless otherwise noted.

## Supplementary figure legends

**Figure S1. Upon feeding following prolonged L1 arrest, *atfs-1(null)* worms develop slower than wildtype worms and are infertile**.

A. Images of wildtype and *atfs-1(null)* worms in control and L1 arrest-fed conditions obtained 4 days after food was introduced (Scale bar, 0.5 mm).

B. Images of wildtype and *atfs-1(null)* worms in L1 arrest-fed conditions obtained at the time wildtype worms reached L4 stage.

**Figure S2. Mitophagy does not impair mtDNA accumulation in *atfs-1(null)* worms following L1 arrest**.

A. Images of wildtype, *atfs-1(null), pdr-1(tm598)* and *atfs-1(null);pdr-1(tm598)* worms in control and L1 arrest-fed conditions obtained when control worms reached the L4 stage (Scale bar, 0.5 mm).

B. Quantification of body length of wildtype, *atfs-1(null), pdr-1(tm598)* and *atfs-1(null);pdr-1(tm598)* in control condition when the worms L4 stage (*n = 3* ± SD, *\*\*\*p* ≤ *0*.*001*, Student’ s t-test).

C. Quantification of body length of L1 arrest-fed wildtype, *atfs-1(null), pdr-1(tm598)* and *atfs-1(null);pdr-1(tm598)* when the control condition worms of each strain reached the L4 stage (*n =* 3 ± SD, *\*\*\*\*p* ≤ *0*.*0001*, Student’ s t-test).

D. mtDNA quantification of control and L1 arrest-fed wildtype, *atfs-1(null), pdr-1(tm598)* and *atfs-1(null);pdr-1(tm598)* worms when the control condition worms of each strain reached the L4 stage (*n = 5* ± SD, *\*\*\*p* = *0*.*0009*, Student’ s t-test).

**Figure S3. Mitochondrial-localized ATFS-1 mediates growth and mitochondrial network recovery following prolonged L1 arrest**.

A. Quantification of TMRE staining morphology of wildtype, *atfs-1(null)* and *atfs-1*^Δ*NLS*^ worms at the L4 stage in control or L1 arrest-fed conditions (*n = 25 to 30 worms*).

B. Images of wildtype, *atfs-1(null)* and *atfs-1*^*R/R*^ worms in control and L1 arrest-fed conditions obtained when control worms reached L4 stage (Scale bar, 0.5 mm).

C. mtDNA quantification of control and L1 arrest-fed wildtype, *atfs-1(null)* and *atfs-1*^*R/R*^ worms when the control condition worms reached the L4 stage (*n = 3* ± SD, *\*p* = *0*.*03* comparing *atfs-1(null)* and *atfs-1*^*R/R*^ worms, *\*p* = *0*.*01* comparing wildtype and *atfs-1(null)* worms Student’ s t-test).

**Figure S4. DAF-2 inhibition promotes mtDNA accumulation and recovery from prolonged L1 arrest**.

A. mtDNA quantification of wildtype and *daf-2(e1370)* starved L1 worms over a period of 5 days (*n = 3* ± SD, *\*p* = *0*.*03* on day 5 comparing wildtype and *daf-2(e1370)*, Student’ s t-test).

B. Images of wildtype and *daf-2(1370)* worms in control and L1 arrest-fed conditions exposed to 0 or 30 μg/ml of ethidium bromide obtained when the control condition worms reached L4 stage (Scale bar, 0.5 mm).

C. mtDNA quantification of wildtype, *daf-2(e1370), daf-16(mu86), daf-2(e1370);daf-16(mu86)* worms at the L4 stage raised in control conditions (*n = 4* ± SD, *\*\*\*p* ≤ *0*.*001*, \**p* ≤ *0*.*05*, Student’ s t-test).

D. mtDNA quantification of wildtype, *daf-2(e1370), daf-2(e1370);daf-16(mu86), daf-2(e1370);atfs-1(null)* and *daf-2(e1370);daf-16(mu86);atfs-1(null)* worms at the L4 stage raised in control conditions (*n = 3* ± SD, *\*\*\*p* ≤ *0*.*001*, \**\*p* ≤ *0*.*01*, Student’ s t-test).

**Table S1.** Data from survival assay of L1s arrested in liquid. Related to figure 1. The following table shows the results obtained for survival assays performed with L1 larvae maintained in solution. The figures the data refer to are indicated in the table. n > 100 worms were used at every time point in every batch.

| Strain | Experiment batch # | Showed in Figure | Temperature (°C) | Mean survival rate | Area under the curve - Alive<sup>##</sup> |
| --- | --- | --- | --- | --- | --- |
| wildtype N2 | 1 | 1C | 20 | $35.26 \pm 0.09$ | 3094 |
| wildtype N2 | 2 | - | 20 | $23.82 \pm 0.14$ | 2116 |
| wildtype N2 | 3 | - | 20 | $33.02 \pm 0.17$ | 2887 |
| *atfs-1(null)* | 1 | 1C | 20 | $34.56 \pm 0.09$ | 2990 |
| *atfs-1(null)* | 2 | - | 20 | $23.52 \pm 0.18$ | 2051 |
| *atfs-1(null)* | 3 | - | 20 | $31.42 \pm 0.21$ | 2724 |

<sup>#</sup> Mean survival rate for each replicate was determined using OASIS2 platform (See also Methods).
<sup>##</sup> Area under the curve was calculated with GraphPad Prism.

**Table S2.** Primer information.

| Strain | Average mean survival rate of all replicates | *p*-value<br>(Student's t-test<br>between the<br>average mean<br>survival rates) |
| --- | --- | --- |
| wildtype N2 | $30.7 \pm 3.5$ | NA |
| *atfs-1(null)* | $29.83 \pm 3.284$ | 0.9364 (n.s) |

**Table S3.** Statistics for lifespan assay of adults. Related to figure 1. The following table shows the results obtained for adult lifespan assay performed on plates. The figures the data refer to are indicated in the table. *n = total 100 to 120* worms for each biological replicate.

| Strain | Experiment batch # | Showed in Figure | Temperature (°C) | Number of worms<sup>#</sup> | Log-rank test $p$-value |
| --- | --- | --- | --- | --- | --- |
| wildtype N2 vs. *atfs-1(null)* | 1 | 1B | 20 | wildtype: 96/100<br>*atfs-1(null)*: 79/100 | ****$p < 0.0001$ |
| wildtype N2 vs. *atfs-1(null)* | 2 | - | 20 | wildtype: 113/120<br>*atfs-1(null)*: 101/120 | ****$p < 0.0001$ |
| wildtype N2 vs. *atfs-1(null)* | 3 | - | 20 | wildtype: 105/110<br>*atfs-1(null)*: 89/110 | ****$p < 0.0001$ |
| wildtype N2 vs. *atfs-1(ΔNLS)* | 1 | 1B | 20 | wildtype: 96/100<br>*atfs-1(ΔNLS)*: 97/100 | n.s |
| wildtype N2 vs. *atfs-1(ΔNLS)* | 2 | - | 20 | wildtype: 113/120<br>*atfs-1(ΔNLS)*: 73/100 | n.s |
| wildtype N2 vs. *atfs-1(ΔNLS)* | 3 | - | 20 | wildtype: 105/110<br>*atfs-1(ΔNLS)*: 96/110 | n.s |

<sup>#</sup>Number of worms represents the number of dead worms scored relative to total number of worms initially started with. The difference in number is indicative of censored worms

