## Supplementary figures for "Mitochondrial genome recovery by ATFS-1 is essential for development following starvation"

**Figure S1:** Upon feeding following prolonged L1 arrest, *atfs-1(null)* worms develop slower than wildtype worms and are infertile

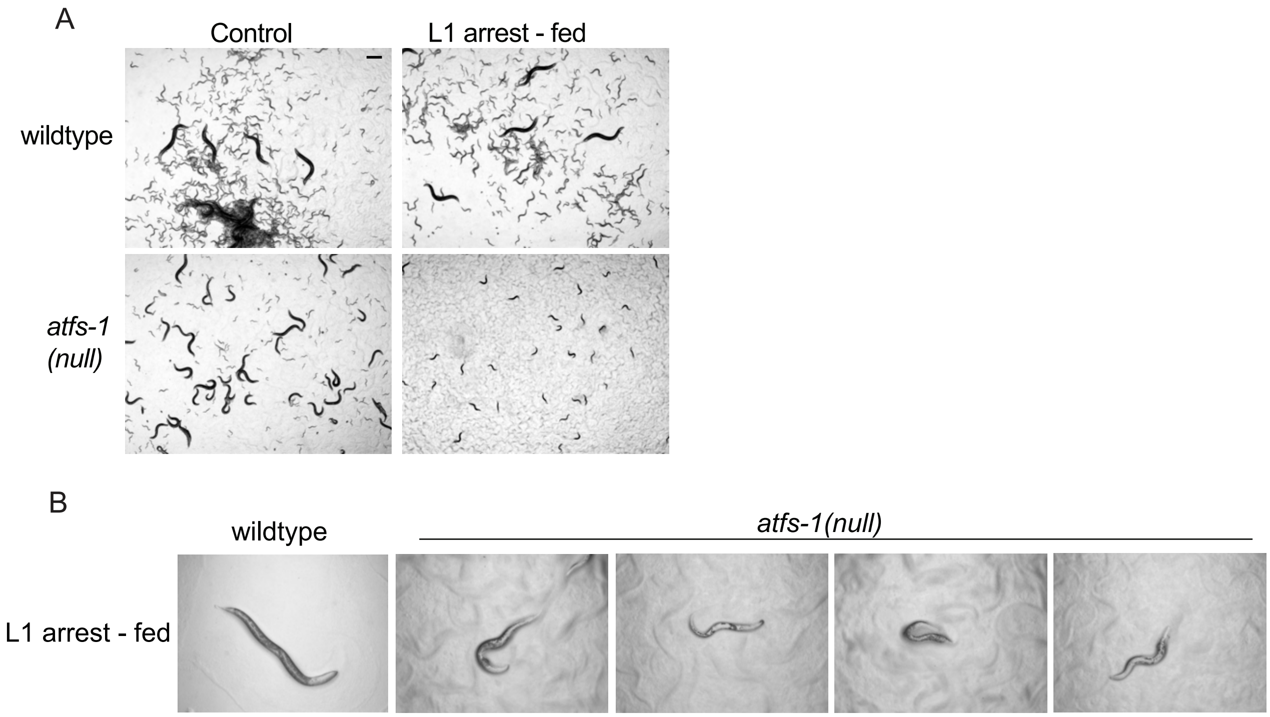

**Figure S2:** Mitophagy does not impair mtDNA accumulation in *atfs-1(null)* worms following L1 arrest

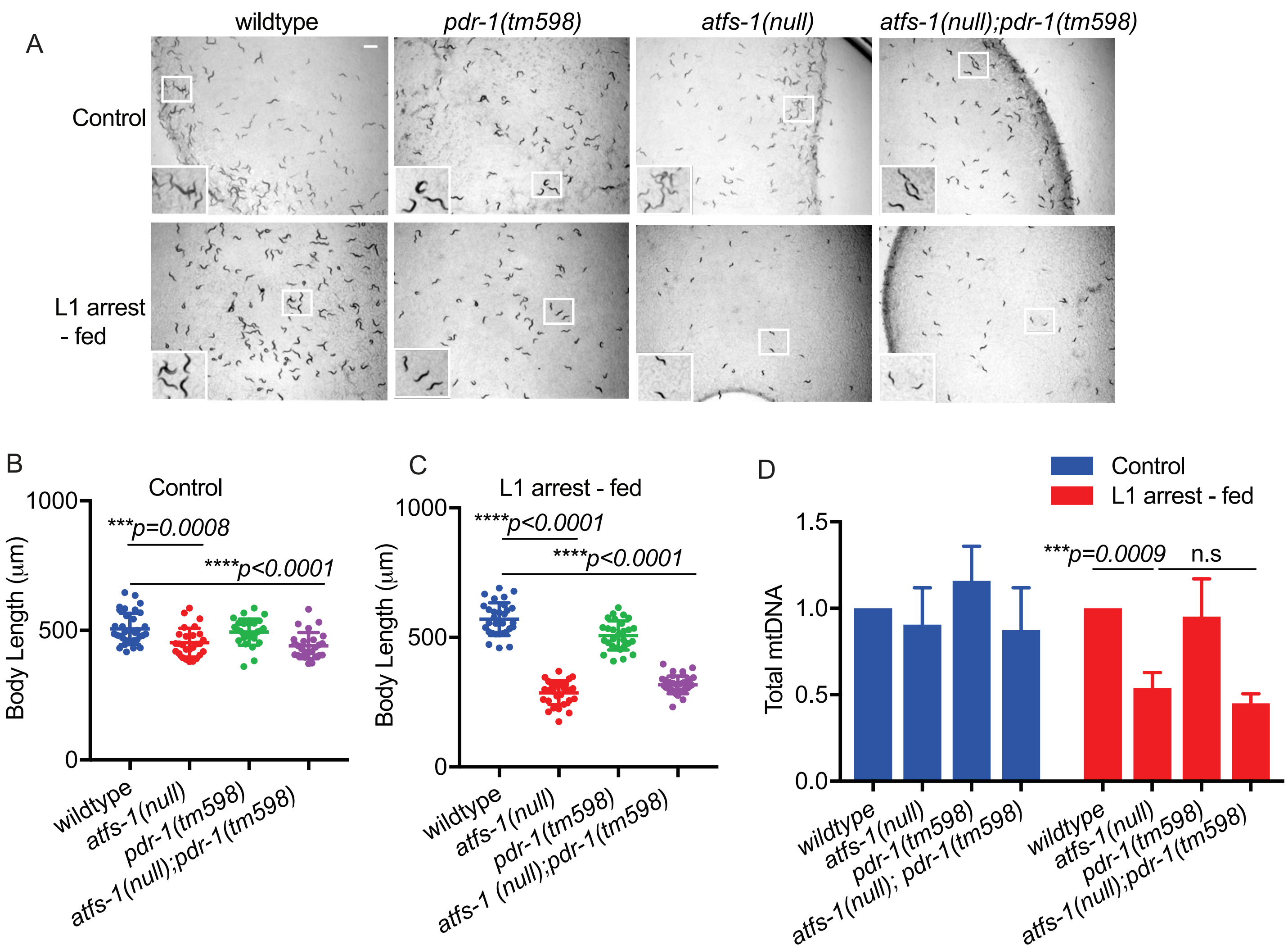

**Figure S3:** Mitochondrial-localized ATFS-1 mediates growth and mitochondrial network recovery following prolonged L1 arrest

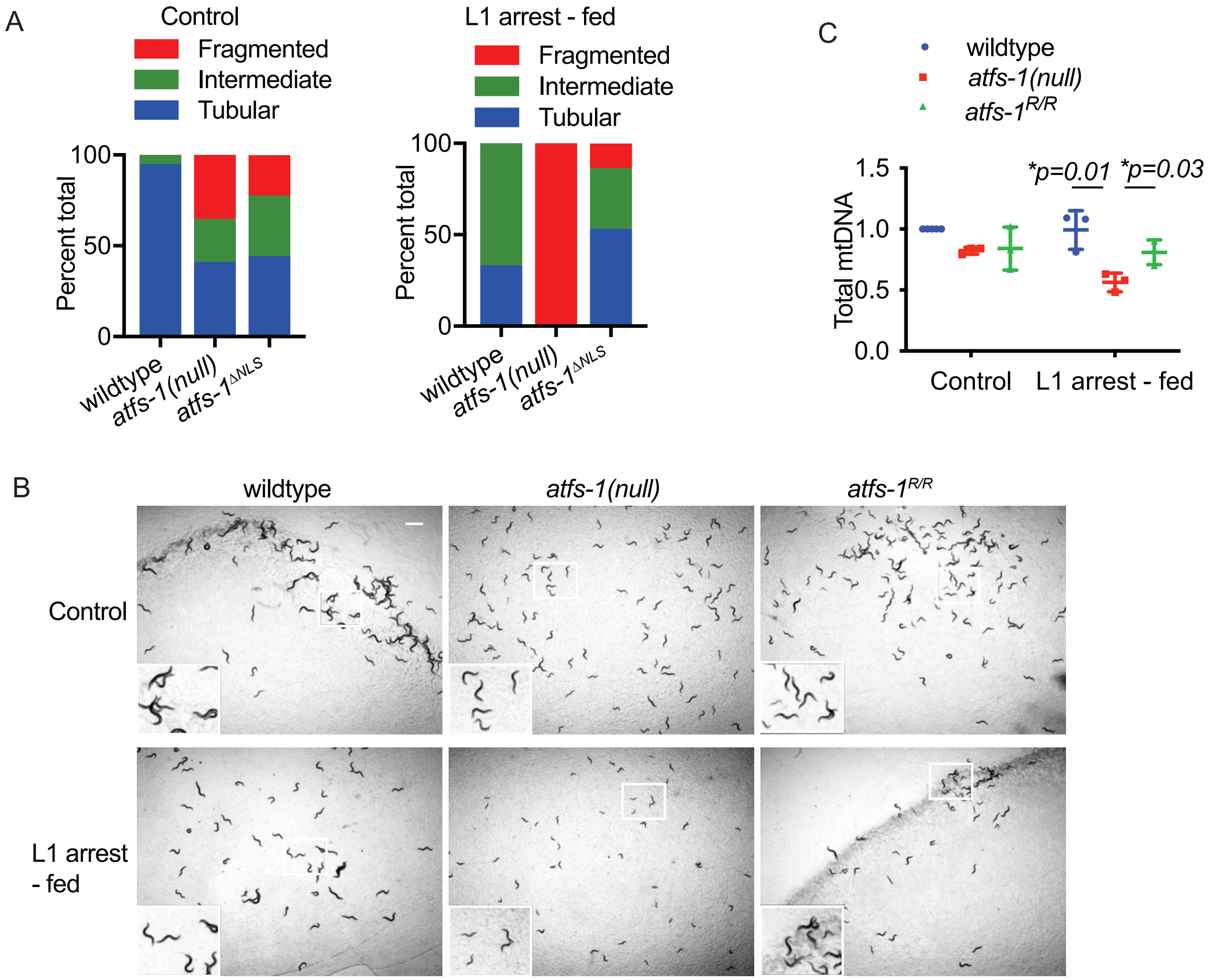

**Figure S4:** DAF-2 inhibition promotes mtDNA accumulation and recovery from prolonged L1 arrest

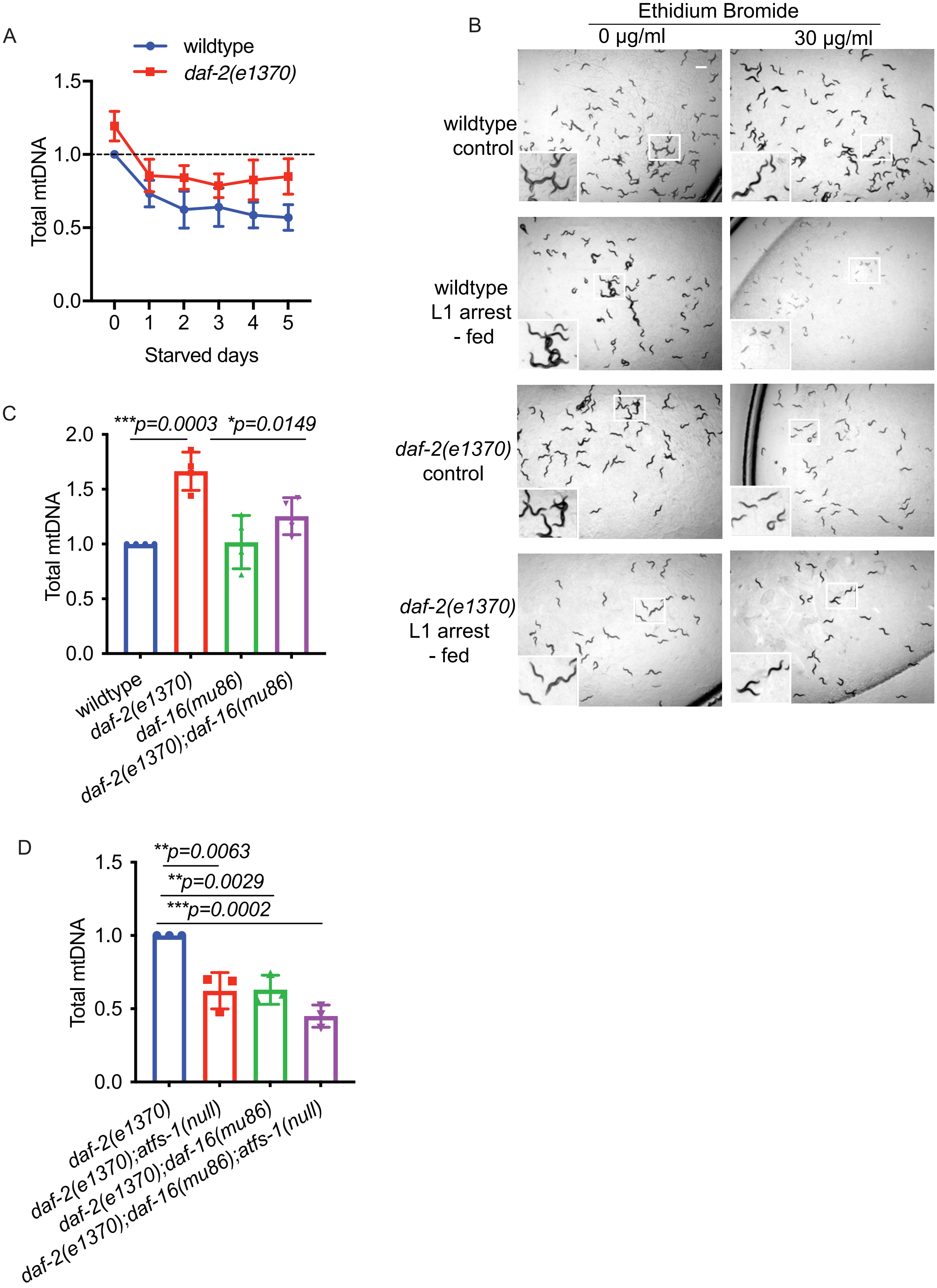
